# MeRIP-seq detects temperature related variation in methylated environmental RNA in fish

**DOI:** 10.1101/2025.11.03.686443

**Authors:** Ehsan Pashay Ahi, Spiros Papakostas, Aristotelis Moulistanos, Tamara Schenekar

## Abstract

Environmental RNA (eRNA) enables non-invasive, near-real-time and potentially functional insight into aquatic communities by capturing RNA molecules released by living organisms. While most eRNA studies focus solely on transcript abundance, this overlooks other regulatory layers that shape how organisms respond to stress. RNA modifications, particularly N^6^-methyladenosine (m^6^A), influence RNA stability, processing, and translation and are known to change under temperature challenges. These properties make m^6^A a compelling target for eRNA-based biomonitoring. We investigated whether m^6^A marks can be recovered from waterborne eRNA released by an African cichlid fish during elevated temperature exposure. eRNA was collected from tank water under control and thermal treatments, enriched for methylated RNA fragments by immunoprecipitation, and sequenced using immunoprecipitation-based MeRIP-Sequencing. Using this approach, we achieved remarkably high read mapping rates to the target fish species, compared to previous studies focusing solely on mRNA sequencing. Moreover, we detect 412 eRNAs having differing abundances between the elevated temperature and control group. Gene ontology analysis reveals that these are enriched in functional categories related to thermal physiology, suggesting a role in the organism’s response to temperature stress. These results indicate an enormous potential of this approach to effectively and non-invasively capture signals of temperature stress response of fish via eRNA. While the current modest replicate depth constrained robust assessment of heat-associated changes in m^6^A, to our knowledge, this is the first attempt to assess m^6^A RNA modification in the context of eRNA research and the first evidence indicating that extra-organismal eRNA can retain epitranscriptomic information.

## Introduction

Non-invasive environmental monitoring based on environmental DNA (eDNA) has matured from concept to practice for tracking biodiversity and ecosystem condition (Veilleux, Misutka, & Glover, 2021)(Blackman et al., 2024). Environmental RNA (eRNA) is now extending this framework: because it is produced by living cells and turns over rapidly, eRNA can deliver a temporally sharper signal than environmental DNA (eDNA), enabling closer-to-real-time biomonitoring of local biological activity (Cristescu, 2019; Kagzi et al., 2023; Wood et al., 2020; Matthew C. Yates, Derry, & Cristescu, 2021). Side-by-side studies show that eRNA often improves concordance with conventional surveys and, paired with eDNA, refines community profiles while reducing false positives linked to legacy signals (Giroux, Reichman, Langknecht, Burgess, & Ho, 2022; Kagzi et al., 2023; Littlefair, Rennie, & Cristescu, 2022; Macher et al., 2024). These features position eRNA as a complement, not a replacement, to eDNA, adding functional and temporal resolution to species detection in environmental samples (Giroux et al., 2022; Littlefair et al., 2022; Matthew C. Yates et al., 2021). Beyond presence-absence, eRNA can deliver information on of organismal physiology without handling target taxa; an urgent need under accelerating warming and more frequent heatwaves (M. C. Yates, Furlan, Thalinger, Yamanaka, & Bernatchez, 2023)(Hechler, Yates, Chain, & Cristescu, 2023). Environmental transcriptomics has detected stress-response transcripts (e.g., heat-shock pathways) directly from extra-organismal eRNA, linking environmental change to molecular function (Hechler et al., 2023). At the same time, the approach faces tractable challenges: RNA of non-microorganismal target taxa is often scarce in an eRNA sample, mapping requires robust reference genomes, and stringent contamination controls are essential (Ahi & Schenekar, 2025; Pochon, Bowers, Zaiko, & Wood, 2025; Wood et al., 2020; Y. Zhang et al., 2024). Emerging study designs and analytical frameworks now address many of these issues, supporting deployment of eRNA for climate-relevant biomonitoring at scale (Bowers et al., 2021; Pochon et al., 2025).

Fishes are particularly valuable sentinels for eRNA-based monitoring: they are ecologically central, economically important, and comparatively well represented in genomic resources (Bowers et al., 2021; Farrell, Whitmore, & Duffy, 2021; Miyata et al., 2021). In aquaria, sequencing of fish-derived eRNA from tank water has captured accumulative stress signatures within hours to days, mirroring organismal condition (Hiki, Watanabe, & Yamamoto, 2024; Miyata, Inoue, Yamane, & Honda, 2025). Field surveys likewise show that fish eRNA metabarcoding can achieve high positive predictivity and, in some cases, outperform eDNA for detecting living assemblages when benchmarked against traditional surveys in rivers and estuaries (Giroux et al., 2022; Littlefair et al., 2022; Miyata et al., 2021). Together, these studies demonstrate that fish eRNA can report both who is present and how populations are responding, information critical for management under rapid environmental change (Stevens & Parsley, 2023).

Importantly, eRNA also contains biochemical context beyond transcript abundance (Ahi & Schenekar, 2025). Post-transcriptional RNA modifications record dynamic regulatory states that shift with environmental stress. N-methyladenosine (m^6^A) is the most abundant internal modification in eukaryotic mRNA and many non-coding RNAs. m^6^A modification is altered via conserved “writer,” “reader,” and “eraser” proteins and it influences splicing, stability, localization, and translation (Zaccara, Ries, & Jaffrey, 2019). Across taxa, including aquatic invertebrates and fishes, temperature stress remodels m^6^A landscapes on stress-responsive transcripts, including heat-shock genes (Ahi & Singh, 2024; Sun et al., 2024; Q. Wang et al., 2025). These observations motivate testing whether epitranscriptomic marks measured in environmental samples can serve as early-warning indicators of physiological stress. Yet, despite rapid growth in environmental transcriptomics, to our knowledge no peer-reviewed study has profiled m^6^A directly from extra-organismal eRNA captured from water while recent roadmaps explicitly call for moving eRNA “beyond mRNA abundance” into the epitranscriptome (Ahi & Schenekar, 2025). Methodological caveats (e.g., antibody specificity and peak-calling reproducibility in immunoprecipitation-based assays) further underscore the need for careful benchmarking (McIntyre et al., 2020).

Here, we address this methodological frontier by applying a conventional m^6^A RNA-sequencing workflow, methylated RNA immunoprecipitation followed by high-throughput sequencing (MeRIP-seq/m^6^A-seq), to water samples containing eRNA shed by the Lake Malawi mbuna cichlid *Labeotropheus trewavasae. L. trewavasae* is well studied and annotated at the transcriptional level and belongs to the haplochromine clade (the most studied group of East African cichlids at transcriptional and genomic levels), which includes closely related sister species with well-annotated genomes, facilitating robust comparative mapping and interpretation (Brawand et al., 2014; Duenser et al., 2023; Roberts, Hu, Albertson, & Kocher, 2011). We expose replicate fish to sub-optimal elevated temperatures in a controlled laboratory setting and enrich collected eRNA for m^6^A-bearing fragments using anti-m^6^A antibodies in a standard MeRIP-seq protocol (Dominissini et al., 2012), enabling transcriptome-wide peak calling. This proof-of-concept establishes feasibility and baseline performance metrics for detecting m A from environmental RNA of a non-model mbuna and lays the groundwork for using environmentally derived epitranscriptomic marks as indicators of thermal stress in *L. trewavasae* and closely related taxa.

## Material and methods

### Experimental design & eRNA sampling

Temperature exposure experiments were conducted in six glass tanks, (volume: 60 L each). Tanks were left fish-free for a minimum of four weeks prior to the experiment and two days prior fish introduction, water in each tank was completely replaced with fresh tap water and the filters and glass surfaces were thoroughly cleaned with tap water to reduce microbial colonization, while maintaining a small microbial biofilm in the filters. Each aquarium was equipped with a submerged filter and oxygenation stone and structural elements (also previously thoroughly cleaned with tap water) were provided for fish cover. To tanks W1-W3 (Table 1), heater rods were added, as well. Nine individuals of *Labeotropheus trewavasae* were introduced (either 4 males and 5 females or 5 males or 4 females, Table 1) to each tank, aiming for an equal total fish biomass of fish among all six tanks (Table 1). At fish introduction, all fish tanks were equilibrated at ambient temperature (26°C). After a 24h acclimation period, heater rods (adjusted to heat up the respective fish tanks to 29°C) were switched on and experimental tanks (W1-W3) equilibrated at target temperature after 12 hours. Temperature of all six fish tanks was carefully monitored, and heater rods were adjusted, if necessary, to stay at the target temperatures (tanks W1-W3: 29°C, tanks C1-C3: 26°C) throughout the entire exposure time span (6 days). After 6 days, eRNA water samples were collected via water filtering through 47mm MCE filters (pore size: 0.45 µm) using a peristaltic pump (MasterFlex LS 600R) connected to a 1 L vacuum flask and 250 ml Nalgene filter holders. Hereby, four replicates per tank were collected and filtering was conducted until filters clogged (24 filters in total, 1.5-3 L per filter, mean: 2.38L, ± 0.54L SD), total water filtered per tank ranged from 6.25L to 12L (Table 1). Filters were transferred into a 5 ml tube immediately after filtration and stored at −80°C until RNA extraction.

**Table 1.** Details on the six fish tanks for temperature exposure experiments of this study.

| **Fish tank** | **Temperature** | **# males** | **# females** | **# fish total** | **biomass total** | **biomass (g/L)** | **Total water filtered (L)** |
| --- | --- | --- | --- | --- | --- | --- | --- |
| W1 | 29°C | 4 | 5 | 9 | 81.27 | 1.35 | 12.00 |
| W2 | 29°C | 5 | 4 | 9 | 81.27 | 1.35 | 12.00 |
| W3 | 29°C | 5 | 4 | 9 | 97.36 | 1.62 | 6.25 |
| C1 | 26°C | 5 | 4 | 9 | 89.32 | 1.49 | 9.50 |
| C2 | 26°C | 5 | 4 | 9 | 89.32 | 1.49 | 8.50 |
| C3 | 26°C | 5 | 4 | 9 | 89.32 | 1.49 | 9.00 |

### RNA preparation & meRIP sequencing

Total RNA was extracted from the filters using Trizol^™^ Reagent (Thermo Fisher Scientific). To increase RNA yield per sample, two sampling replicates per tank were pooled at the final step of the extraction (elution of the RNA pellet), leaving two independent eRNA samples per tank (12 samples in total) and a final elution volume of 30 µl per pooled sample. RNA extracts were subjected to DNase I digestion (New England Biolabs) followed by rRNA depletion using the NEBNext rRNA depletion Kit (New England Biolabs) according to the manufacturer’s instructions. rRNA depleted samples were fragmented with the NEBNext Magnesium RNA Fragmention Module (New England Biolabs) using a 5 min incubation time. Fragmented RNA was enriched for N^6^-methyladenosine using the EpiMark® N6-Methyladenosine Enrichment Kit (New England Biolabs) according to the manufacturer’s instructions.

Enriched samples were sent to Novogene for RIP-Seq library preparation and sequencing and sequenced via illumina sequencing in PE150 read mode with a targeted output of 79G per sample.

### Bioinformatic analysis

Raw Illumina paired-end reads were first subjected to quality control using FastQC (Andrews, 2012), to assess base quality, adaptor content, and sequence duplication levels. Adapter trimming and quality filtering were then performed using Trimmomatic v0.39 (Bolger, Lohse, & Usadel, 2014) with following parameters: we used ILLUMINACLIP to remove Illumina adapter sequences allowing up to two mismatches, a palindrome clip threshold of 30, and a simple clip threshold of 10; LEADING and TRAILING parameters to trim bases with Phred quality scores below 5 at the start and end of reads, respectively; SLIDINGWINDOW:3:20 to trim once the average Phred score within a 3-bp window drops below 20; MINLEN:100 to discard reads shorter than 100 bp after trimming. The resulting high-quality paired reads were screened and mapped against the target species *Labeotropheus trewavasae* (NCBI Assembly ASM4766397v1) using FastQ Screen (Wingett & Andrews, 2018) with Bowtie2 (Langmead & Salzberg, 2012) as the alignment engine. Fractions of mapped reads were compared between control and treatment groups using Wilcoxon rank-sum test and potential correlations between the fractions of mapped reads and filtered water volume, as well as biomass were assessed via Spearman’s rank correlation tests. Mapped reads were tagged and extracted with a custom Python-script subsequently used to reassemble validated mate-pairs. To facilitate downstream functional analyses, the extracted reads were re-mapped against the phylogenetically related and well-annotated model genome of *Oreochromis niloticus* (NCBI Assembly GCF_018398535.1) using the BWA-MEM algorithm (H. Li & Durbin, 2009) in paired-end mode. Properly paired and uniquely mapped reads were retained using SAMtools and sra-tools for subsequent analyses. Differential gene expression between high temperature and control samples was assessed using metaseqR2 (Fanidis & Moulos, 2021; Moulos & Hatzis, 2015), with Benjamini–Hochberg adjustment to control the false discovery rate; genes with padj < 0.05 were considered significant. Condition-specific transcriptional profiles were analyzed using Principal Component Analysis (PCA) with the R function prcomp. Hierarchical clustering was performed and visualized using pheatmap. Significant genes were further represented in a volcano plot, generated with ggplot2 and plotly. All analyses were conducted in R v.4.4.1 (R Core Team, 2023).

## Results

### Mapping of MeRIP-eRNA reads to *L. trewavasae*

On average, 581,909,399 (± 149,038,971 SD) raw reads were produced per sample. Of the samples passing quality filtering steps, an average of 22.6% (± 5.3 SD) of the reads mapped to the L. trewavasae reference at 99% identity (Table 2), across all twelve MeRIP-eRNA libraries (two per tank), with values spanning 13.6–29.4%. Mapping fractions were broadly similar between temperature conditions: the warmer tanks (29 °C; W1–W3) averaged 24.2% (± 5.9), while the control tanks (26 °C; C1–C3) averaged 21.0% (± 4.0), and an exact Wilcoxon rank-sum test indicated no statistically significant difference between treatments (W = 11, p = 0.31). Within the warmer condition, W1 and W2 showed generally higher values (W1: 28.6–29.3%; W2: 26.0–29.4%), whereas W3 was uniformly lower (15.4–16.8%); controls were intermediate to high (C1: 20.5–21.4%; C2: 13.6–20.1%; C3: 23.8–26.6%). Group medians were 27.3% for the warmer condition and 21.0% for controls (overall median 22.6%) (Table 2).

**Table 2:** Percentages of processed reads that could be mapped to *L. trewavasae* with 99% similarity. For each sample details of the sample origin, such as fish tank and exposed temperature are shown. Percentages of each sample represent mean value across forward and reverse reads. Furthermore, mean / median values together with standard deviation (±) for the two temperature groups, as well as total overall are given.

| eRNA sample | fish tank | temperature | % reads mapped | biomass (g/L) | water filtered (L) |
| --- | --- | --- | --- | --- | --- |
| W1_1 | W1 | 29°C | 28.6 | 1.35 | 6 |
| W1_2 | W1 | 29°C | 29.3 | 1.35 | 6 |
| W2_1 | W2 | 29°C | 29.4 | 1.35 | 6 |
| W2_2 | W2 | 29°C | 26.0 | 1.35 | 6 |
| W3_1 | W3 | 29°C | 16.8 | 1.62 | 3.25 |
| W3_2 | W3 | 29°C | 15.4 | 1.62 | 3 |
| C1_1 | C1 | 26°C | 20.5 | 1.49 | 5 |
| C1_2 | C1 | 26°C | 21.4 | 1.49 | 4.5 |
| C2_1 | C2 | 26°C | 13.6 | 1.49 | 4.75 |
| C2_2 | C2 | 26°C | 20.1 | 1.49 | 3.75 |
| C3_1 | C3 | 26°C | 23.8 | 1.49 | 5 |
| C3_2 | C3 | 26°C | 26.6 | 1.49 | 4 |
| Overall group W | | 29°C | 24.2 / 27.3(± 5.9) | 1.44 / 1.35 (± 0.13) | 5.04 / 6.00 (± 1.36) |
| Overall group C | | 26°C | 21.0 / 21.0 (± 4.0) | 1.49 / 1.49 (± 0) | 4.50 / 4.63 (± 0.48) |
| Overall total | | - | 22.6 / 22.6(± 4.0) | 1.47 / 1.49 (± 0.09) | 4.77 / 4.88 (± 1.05) |

### Mapping rates vs. water volume and biomass

Across all twelve eRNA libraries, the fraction of processed reads mapping to L. trewavasae increased with the total volume of water filtered and decreased with fish biomass in the tank (Fig. 1). Spearman’s tests showed a positive association with water filtered (ρ = 0.752, S = 70.8, p = 0.005) and a negative association with biomass (ρ = −0.817, S = 519.6, p = 0.001). These trends are also observable through sloping linear regression lines in Figure 1.

**Fig. 1:**
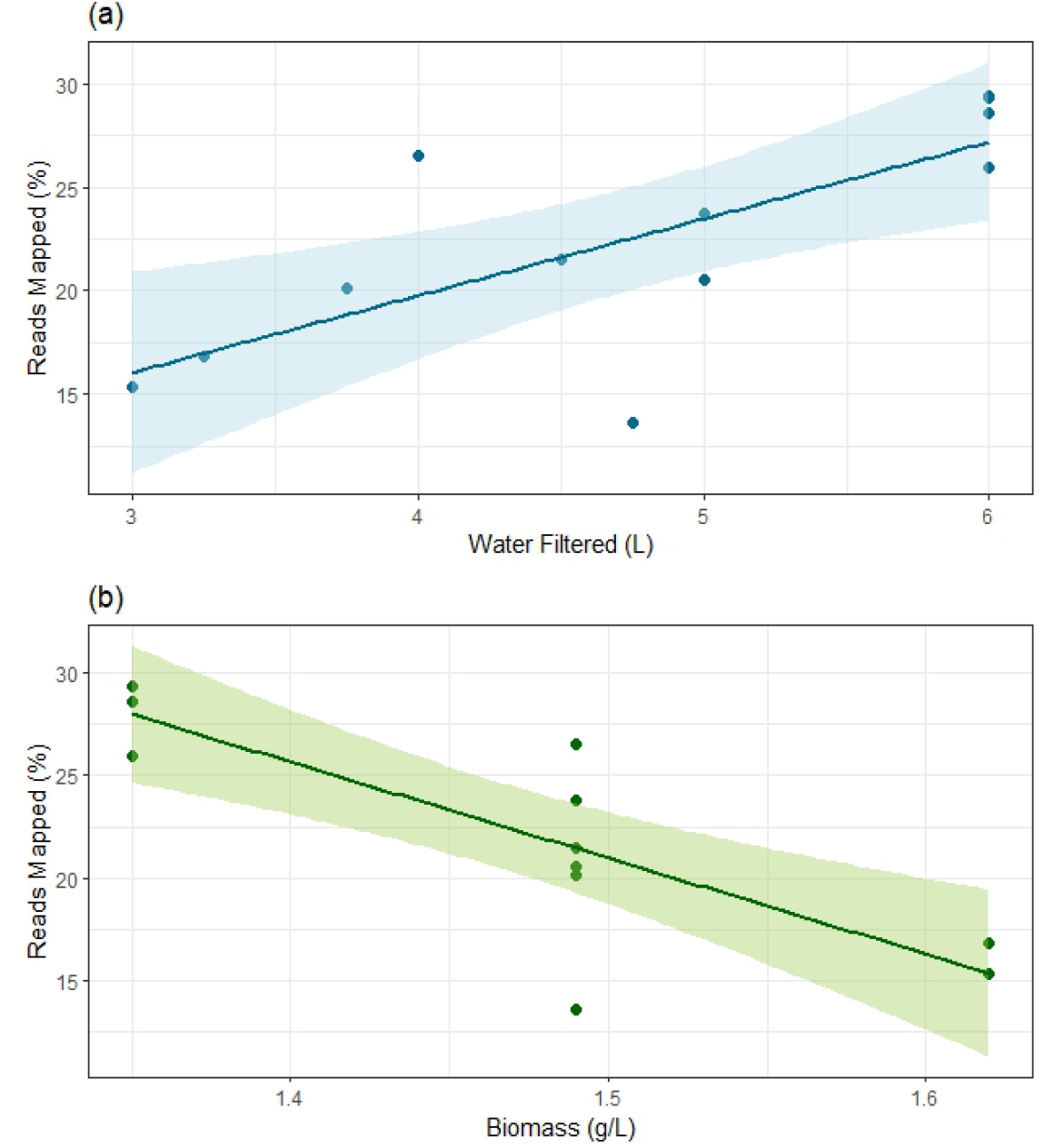
Relationship between reads mapped (%) and water volumes filtered, as well as fish biomass in the experimental tanks across all eRNA samples analyzed. **(a)** Scatter plot of reads mapped (%) versus water volumes filtered, showing a positive correlation. **(b)** Scatter plot of reads mapped (%) versus fish biomass, showing a negative correlation. Points represent individual data points, and lines represent linear regression trends with 95% confidence intervals (shaded areas).

**Fig. 2:**
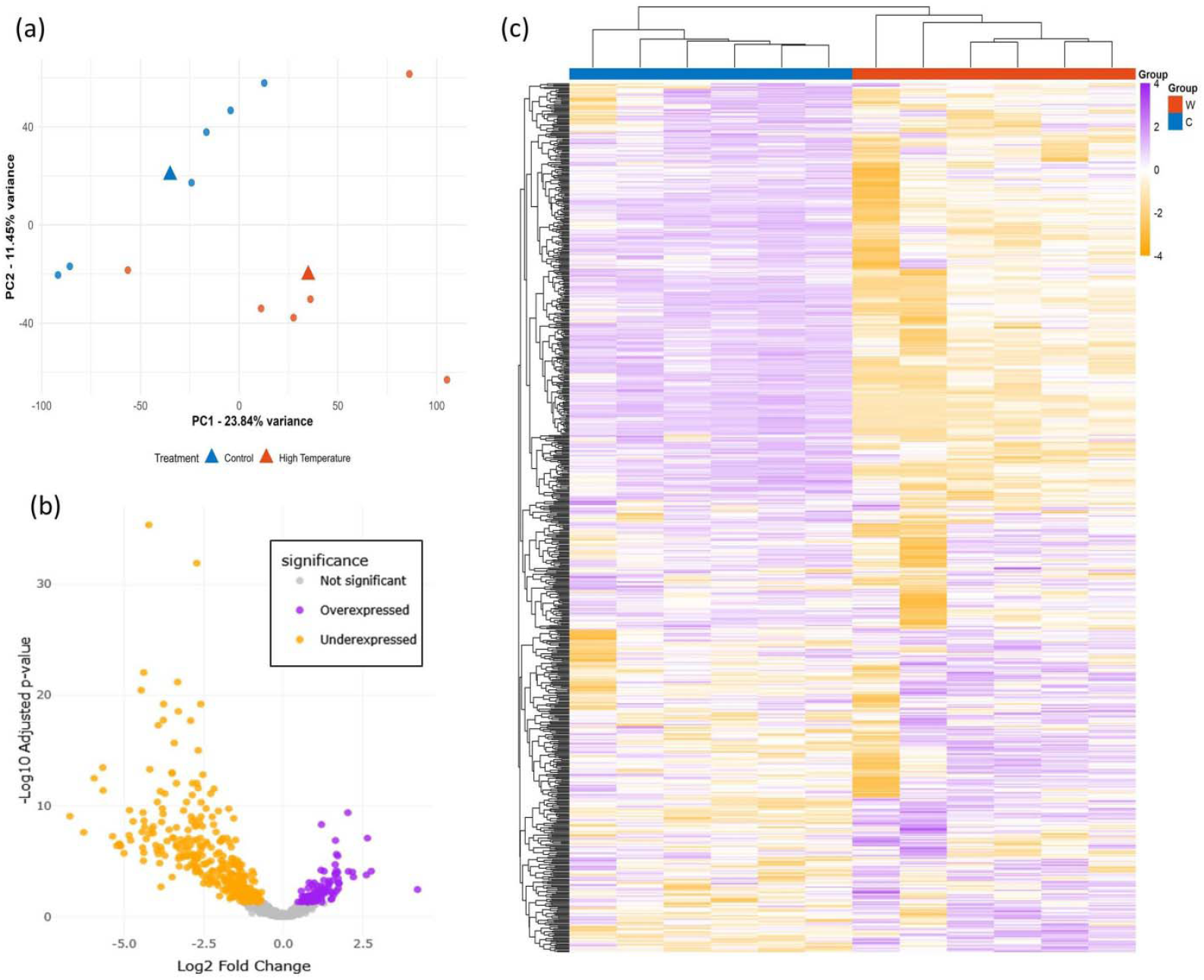
Differential abundance of m6A-enriched eRNAs in warm vs. control tanks. **(a)** PCA of transcriptomic profiles of the quantified 755 transcripts under control and high-temperature conditions. Circles represent samples colored by treatment (Control = blue, High Temperature = orange). Triangles indicate group centroids. Axes show PC1 and PC2 with the percentage of variance explained. **(b)** Volcano plot of differential eRNA abundances between high temperature and control conditions. Each point represents a transcript (total n = 755). Significant transcripts are shown in purple and orange (padj < 0.05) and non-significant genes in gray. The x-axis represents log2 fold change, and the y-axis represents −log10 adjusted p-value. **(c)** Heatmap of gene expression in Control and High Temperature samples. Heatmap shows row-scaled (Z-score) log2-transformed expression values for 755 quantified transcripts across 12 samples. Purple indicates higher relative expression, white intermediate, and orange lower expression. The colored bar above columns indicates treatment: Control (blue) and High Temperature (orange). Genes and samples were clustered using hierarchical clustering (Euclidean distance, complete linkage).

### eRNAs more abundant in warmer tanks

Out of 755 detected eRNAs, 106 showed significantly higher abundance in warmer tanks (FDR < 0.05; 14.0% of all detected), with log2-normalized fold changes ranging from 0.467 to 4.213 (median 1.085; mean 1.207), corresponding to ∼1.38×–18.55× higher abundance (median ∼2.12×); 62/106 (58.5%) were ≥2× (log2FC ≥ 1), 26/106 (24.5%) were ≥∼2.83× (log2FC ≥ 1.5), and 8/106 (7.5%) were ≥4× (log2FC ≥ 2), with the most statistically significant characterized transcripts being *klf4, abca12, capn9, lmo7a, cdc42bpb, dsg2l, mid1ip1a, myh14, znf395b*, and *znf185*, and the largest characterized fold changes observed for *mov10l1, capn9, btr12, arhgap44, znf395b, paqr8, klf4, stk10, pdlim2*, and pard6a; by annotation category, 82/106 (77.4%) corresponded to protein-coding–like mRNAs and 24/106 (22.6%) to uncharacterized loci/clone designations. GO enrichment on these 106 genes identified three significant terms: intermediate filament-based process (GO:0045103; FDR = 0.0021), antigen processing and presentation (GO:0019882; FDR = 0.032), and cell–cell adhesion (GO:0098609; FDR = 0.045), consistent with cytoskeletal remodeling, epithelial barrier/junction dynamics, and immune interface changes under warmer conditions.

### eRNAs less abundant in warmer tanks

A total of 306 transcripts showed significantly lower abundance in warmer tanks (FDR < 0.05), with log2-normalized fold changes ranging from −0.689 to −6.700 (median −2.069; mean −2.359), corresponding to ∼1.61×–104× lower abundance (median ∼4.20×; mean ∼5.13×); 279/306 (91.2%) were ≥2× lower (log2FC ≤ −1), 212/306 (69.3%) were ≥∼2.83× lower (log2FC ≤ −1.5), and 158/306 (51.6%) were ≥4× lower (log2FC ≤ −2). The most statistically significant characterized transcripts included *mmp9, cebpb, antxr2a, abr, dennd4a, f8, marcksl1a, socs3a, tktb*, and *bcl2l1*; the largest characterized fold changes were observed for *marcksl1b, ldha, eno1a, sh2d3cb, adam28, rock2a, lrrc23, sytl3, dennd4a*, and *kcnj2a*. By annotation category, 218/306 (71.2%) corresponded to protein-coding–like mRNAs and 88/306 (28.8%) to uncharacterized loci/clone designations. GO enrichment on these down-regulated genes identified three significant terms: pyridine-containing compound metabolic process (GO:0072524; FDR = 2.16 × 10^?4^), small GTPase-mediated signal transduction (GO:0007264; FDR = 0.0237), and purine-containing compound metabolic process (GO:0072521; FDR = 0.0415), consistent with a coordinated metabolic downshift (notably glycolysis/PPP and nucleotide/NAD(H) pathways) and dampened GTPase-dependent signaling/cytoskeletal programs under warmer conditions.

## Discussion

Environmental RNA (eRNA) offers the potential for non-invasive aquatic monitoring, often confirming conventional species assemblage survey methods while adding temporal resolution (Giroux et al., 2022; Littlefair et al., 2022; Miyata et al., 2021, 2025; Veilleux et al., 2021; M. C. Yates et al., 2023; Matthew C. Yates et al., 2021). However, eRNA’s true potential lies beyond presence-absence monitoring but in its ability to convey information about physiological conditions of organisms (Ahi & Schenekar, 2025; M. C. Yates et al., 2023). Beyond transcript counts, chemical marks on RNA, especially m^6^A, shape RNA stability, processing, and translation, and they shift with temperature in diverse aquatic taxa (Ahi & Singh, 2024; Sun et al., 2024; Q. Wang et al., 2025). Recent overviews make clear that mRNA modifications are moving into applied contexts, including environmental biology (Ahi & Schenekar, 2025). Against that backdrop, our study asked two first-order questions: (1) can extra-organismal eRNA from tank water be taken through a conventional antibody-based workflow to obtain material suitable for m^6^A profiling in a cichlid? and (2) Can this m^6^A – modified eRNA be utilized to gain information about environmental cues, such as environmental stressors promoting physiological responses organisms?

All MeRIP-eRNA libraries contained target fish RNA signals, and mapping percentages were similar between the warmer (29 °C) and control (26 °C) conditions (Table 2). Values fell within a practical range for downstream work (∼23% on average at 99% identity), replicate libraries within tanks were closely matched, and one warmer tank consistently mapped lower than its peers. These results indicate that the fraction of host-mapped reads did not shift between treatments in this design and that library quality is sufficient to proceed to functional analyses. These mapping rates are two orders of magnitude higher of those achieved by previous eRNA studies assessing stress response in macro-organisms but focusing on conventional mRNA-Seq (Hechler et al., 2023; Hiki, Yamagishi, & Yamamoto, 2023). The higher mapping rates could potentially result from the higher prevalence of m^6^A modifications in eukaryotic compared to prokaryotic transcripts and could suggest m^6^A – targeted eRNA analysis as a solution for the reported microbial RNA overload in metatranscriptomic analysis.

Observed correlations point to two practical drivers of target species-mapping yield: (1) filtered water volume and (2) biomass effects. For the first one, larger filtered water volumes likely improved capture of rare target taxon eRNA fragments, an expected effect reported also from eDNA work, (Bowers et al., 2021; Capo, Spong, Königsson, & Byström, 2020; Hinlo, Gleeson, Lintermans, & Furlan, 2017; Spens et al., 2017). Conversely, tanks with greater biomass showed lower host-mapping fractions. While we expected a reverse pattern, this could potentially result from higher organic input through defecation in higher densities, stimulating microbial growth. This could have resulted in a higher non-target-microbial to target-fish eRNA ratios, and the microbial reads taking up a higher fraction of the produced sequencing reads (Adiconis et al., 2013; Gann, Kang, Dyhrman, Gobler, & Wilhelm, 2021; Gifford, Sharma, Rinta-Kanto, & Moran, 2011; Herbert et al., 2018; Pereira-Marques, Ferreira, & Figueiredo, 2024; Wood et al., 2020; Xia et al., 2024). In practical terms, the patterns in Fig. 1 argue for standardizing filtered volume across treatments in similar experiments, logging biomass (or a correlated metric) as a model covariate, and keeping library preparation treatments tightly consistent. (Herbert et al., 2018; Xia et al., 2024).

### Thermal stress elevated m^6^A-enriched eRNAs and their functional relevance

Among the detected transcripts, 106 genes had higher abundance in warmer tanks, and their enrichment for intermediate filament–based process (GO:0045103), cell–cell adhesion (GO:0098609), and antigen processing and presentation (GO:0019882) aligns with established vertebrate responses to thermal challenge in barrier epithelia. Heat elevates epithelial permeability and disrupts tight-junction architecture in intestine and other exposed surfaces, placing adhesion machinery at the center of hyperthermia-induced barrier failure (Dokladny, Zuhl, & Moseley, 2016; Patra & Kar, 2021; Toivola, Strnad, Habtezion, & Omary, 2010). In parallel, keratin intermediate filaments are classic heat-responsive scaffolds that remodel and undergo phosphorylation during heat shock to preserve mechanical integrity (Kayser et al., 2013; Toivola et al., 2010). Heat-shock programs can also intersect antigen handling (e.g., HSP-assisted cross-presentation), providing a plausible link to the immune GO term under warming (Murshid, Gong, & Calderwood, 2012).

Within this framework, several high-signal candidates map to barrier modules that are both thermally implicated and expressed in water-exposed tissues (skin, gill, gut). The tight-junction scaffold *tjp1a* (zo-1) sits at the core of heat-induced intestinal leakiness (Dokladny et al., 2016; Patra & Kar, 2021; Toivola et al., 2010). The junctional organizer *lmo7a* strengthens apical actomyosin by bridging nectin–afadin and e-cadherin–catenin systems and recruiting non-muscle myosin II—functions central to epithelial mechanics at exposed surfaces (Fu, Jiang, Li, Zhu, & Zhang, 2022; Matsuda, Chu, & Sokol, 2022; Ooshio et al., 2004). Desmosomal components (*dsg2l, dsp(a/b), pkp3a*) anchor keratins to the membrane to resist mechanical load, supporting barrier integrity when temperature rises (Fülle et al., 2024; Gross et al., 2018; Johnson, Najor, & Green, 2014). On the intermediate-filament axis, *krt8* (and epidermal *krt15*) are directly heat responsive, with keratin networks reorganizing under heat shock; cytolinkers such as *evpla* and *ppl* stabilize these networks and are abundant in skin and mucosa (Kayser et al., 2013; Toivola et al., 2010). Additional top signals with thermal literature include the transcription factor *klf4*, induced rapidly by heat stress in vivo and in cells (Liu et al., 2006); *dusp5*, a nuclear ERK phosphatase induced by heat shock (Seo et al., 2017); *foxo3b* (ortholog of FOXO3) under an HSF1→FOXO3 axis activated by heat shock (Grossi et al., 2018)(Bernardo, Torres, & da Silva, 2023); slc1a5, upregulated after heat stress in cattle (Dado-Senn, Skibiel, Dahl, Apelo, & Laporta, 2021); and *mcu*, whose expression increases in immune organs of heat-stressed broilers, consistent with heat-linked Ca^2+^/mitochondrial remodeling (Yong Wang et al., 2023). Finally, the barrier-lipid transporter *abca12*, essential for lamellar-body cargoing and epidermal permeability, supports a skin-source interpretation for eRNA despite not being a canonical heat marker (Akiyama, 2014; Leprince & Simon, 2025). Taken together, these patterns ground the GO results in thermal biology: junctional and IF programs that protect (or remodel) the epithelial interface under warming, and immune presentation consistent with heat-evoked epithelial stress.

Given that these eRNAs were captured by meRIP-seq, it is notable that several of the most interesting transcripts also carry independent m^6^A evidence in vertebrates: *klf4* (MeRIP-qPCR–validated m^6^A in intestinal epithelium) (Zhu et al., 2024); *foxo3b/foxo3* (YTHDF3-mediated translation via m^6^A sites) (Hao et al., 2022); *slc1a5* (stabilization via the IGF2BP2–m^6^A axis) (Pu et al., 2025); junctional/adhesion factors (*lmo7, znf185*) among differentially m^6^A-modified genes in MeRIP-seq datasets (C. Lin et al., 2023); paqr8 identified as highly m^6^A-methylated at the isoform level in human brain maps (Gleeson et al., 2025); and *mov10l1* supported by the mammalian *MOV10* ortholog functioning as an m^6^A reader that promotes decay of m^6^A-bearing mRNAs (Mehravar et al., 2022). These orthogonal reports reinforce an epitranscriptomic layer acting on key barrier and signaling transcripts that also rise in abundance in the warmer tanks.

### Thermal stress reduced m^6^A-enriched eRNAs and their functional relevance

The GO signals in our down-regulated set; pyridine- and purine-containing compound metabolism plus small GTPase–mediated signaling, fit known axes of thermal physiology in fishes that live behind water-exposed epithelia. Heat elevates oxidative and osmotic loads in gill and gut, with histology and transcriptomics repeatedly showing epithelial damage, barrier dysfunction and redox strain under warming. This places a premium on NADPH economy and nucleotide turnover for repair and antioxidant buffering (S. Yang et al., 2023)(Stincone et al., 2015)(Madeira, Vinagre, & Diniz, 2016)(Zhou, Gao, & Wang, 2023) (Ahi, Lindeza, Miettinen, & Primmer, 2025). In that context, the prominence of glycolysis/PPP genes (e.g., *ldha, pgd, tktb, taldo1*) among our top-ranked warm-decreased eRNAs is consistent with an acclimatory set-point in which basal glycolytic throughput and PPP-supplied NADPH are dialed back after the acute phase, a pattern echoed in experimental work tracking metabolic reprogramming during sustained heat exposure (Stincone et al., 2015)(Madeira et al., 2016)(H. Qin et al., 2023). Thermal injury also forces rapid remodeling at exposed surfaces, and cytoskeletal/adhesion programs are a recurring feature of heat responses in fish epithelia. In gill, heat-tolerant and heat-sensitive lineages differ strongly in extracellular matrix, junction and adhesion signatures after heat challenge; in intestine, warm exposure disrupts tight-junction composition and increases permeability, with focal adhesion and actin regulation among the enriched pathways (Asakawa et al., 2019; Zhou et al., 2023). The down-ranked small-GTPase/cytoskeletal cohort in our data (including *rock2a* and regulators of Rho/Arf/Ras signaling) fits well with these observations and suggests a longer-term state that reduces high-cost cytoskeletal turnover in warm water, adaptive for energy budgeting, but potentially limiting barrier resealing when damage accumulates (Asakawa et al., 2019; Zhou et al., 2023) (Ahi et al., 2025).

Among individual candidates, several have direct thermal evidence and clear relevance to water-exposed tissues. The *nod2–ripk2* axis shows temperature sensitivity in cyprinids: heat stress induces nod2 and its adaptor *rick/ripk2* across gill, liver, kidney and blood, indicating thermal engagement of cytosolic pattern-recognition pathways at mucosal interfaces (Basu et al., 2015). Given that gill and intestine routinely suffer heat-induced damage and dysbiosis, a chronic reduction of *nod2/ripk2* eRNAs under warm tanks may reflect a post-acute resetting of innate tone to limit costs of constant surveillance at the interface (S. Yang et al., 2023) (Madeira et al., 2016) (Zhou et al., 2023). The cytokine brake socs3 is also heat-responsive in teleosts, upregulated within hours during acute warming in Chinese tongue sole brain and altered under chronic temperature regimes in gene-panel assays, supporting a role for JAK/STAT damping in thermal acclimation; its lower eRNA abundance here aligns with a longer-term return toward baseline signaling (Yue Wang et al., 2024).

A second cluster centers on matrix turnover and epithelial mechanics. mmp9 is expressed in teleost mucosae (including gill and skin) and orchestrates extracellular-matrix remodeling during inflammation and repair; thermal acclimation studies have linked warm exposure to changes in MMP-associated remodeling, and independent work shows that heat damages gill epithelia, the very condition in which MMP-dependent turnover is typically engaged (Chadzinska, Baginski, Kolaczkowska, Savelkoul, & Lidy Verburg-Van Kemenade, 2008)(Keen, Mackrill, Gardner, & Shiels, 2021) (S. Yang et al., 2023). The coordinated warm-decrease of *mmp9* (with other protease/antiprotease partners in our list) is therefore compatible with an energy-saving, low-turnover state at the gill surface, albeit with the trade-off of slower matrix renewal under sustained heat (S. Yang et al., 2023) (Keen et al., 2021).

Thermal links are also strong on the metabolic side. Across species, heat challenges often boost lactate/LDH during the acute window, tracking a transient reliance on anaerobic glycolysis (H. Qin et al., 2023); our warm-decreased *ldha* and allied glycolysis/PPP eRNAs thus look like the opposite, acclimatory phase, an energetically conservative setting once immediate heat shock has passed, consistent with the broader literature on metabolic and osmotic re-set in gills under warming (S. Yang et al., 2023)(Madeira et al., 2016)(H. Qin et al., 2023). This model is also compatible with temperature–oxygen cross-talk involving *hif1a/hif1aa* targets, which frequently intersect with heat responses in fishes, even if the contribution of HIF-1 to short-term heat tolerance can be context-specific (Madeira et al., 2016).

Finally, several of our priority transcripts carry direct m^6^A marks in vertebrates, offering a plausible post-transcriptional mechanism for the persistent down-shift we observe under warm tanks. m^6^A deposition enhances *ldha* translation via YTHDF1 and helps sustain glycolysis; eno1 translation is likewise promoted by m^6^A readers; *slc2a1/glut1* is stabilized via METTL3–IGF2BP2/3; *hif-1α* translation is supported by m^6^A/YTHDF1 during hypoxic demand; and both *mmp9* and *socs3* have been shown to be m^6^A-regulated (Chen, Wu, Zuo, Wu, & Chen, 2025; Q. Li et al., 2021; Z. Li, Meng, Chen, Xu, & Guo, 2023; Ma et al., 2022; Shen et al., 2020; K. Zhang et al., 2022). Given the ranking and tissue relevance of *ldha, eno1a, slc2a1b, hif1aa, mmp9* and *socs3a* in our dataset, conserved m^6^A pathways are compelling candidates for coupling sustained warmth to altered RNA stability/translation at gill and gut surfaces.

### Methodological and conceptual aspects for future m^6^A-targeted eRNA research

Practical limits in environmental transcriptomics exist, such as scarce target RNA, microbial and rRNA overload and degraded target fragments and can sway usable RNA yield and reproducibility (Bowers et al., 2021; Wood et al., 2020). Thus, rRNA depletion chemistry matters. cross-site tests show that kits differ in efficiency with fragmented inputs (Herbert et al., 2018). Because m^6^A modifies multiple RNA classes, not only mRNA but also many other types of RNAs such as circRNAs and miRNAs (Mei et al., 2023; S. Qin et al., 2022), which can be more stable than mRNAs, a modification-centric readout may retain relevant environmental signal when long mRNA fragments are scarce in eRNA samples (Ahi & Schenekar, 2025; Köberle et al., 2013; Y. Li et al., 2015; T. Lin & Meegaskumbura, 2025; Quan et al., 2021; Silva, Lopes, Teixeira, Sousa, & Medeiros, 2015; S. Wang et al., 2021). In addition, vesicle-protected small RNAs can persist outside cells, and because demethylation is enzyme-dependent (FTO/ALKBH5) rather than spontaneous, the N^6^-methyl mark is expected to persist as long as the RNA remains intact (Arroyo et al., 2011; Gao, Zha, Li, Xia, & Wang, 2024; Pan et al., 2021). However, concerns specific to m^6^A IP assays (antibody specificity, peak reproducibility) require careful attentions when starting from low abundant extra-organismal eRNA (McIntyre et al., 2020). There is precedence that modification states are measurable in RNA released outside cells: modified nucleosides, including m^6^A, have been quantified in exosomal small RNAs and in serum, and m^6^A on miRNAs can bias their export into extracellular vesicles (Garbo et al., 2024; Guo, Hu, Cao, & Wang, 2021; Hu et al., 2021; Pan et al., 2021). Such findings suggest that changes in modification patterns are detectable in extracellular material, supporting our initial plan to analyze m^6^A peaks directly in eRNA.

Finally, several approaches are available for very low-abundance RNA if subsequent iterations require more sensitivity or resolution. Antibody-independent or locus-tuned methods (DART-seq, MAZTER-seq, m^6^A-SAC-seq) and optimized antibody-based or direct RNA nanopore strategies (miCLIP2, m^6^A calling from nanopore current) reduce input needs or deliver base-level information (Garcia-Campos et al., 2019; Ge et al., 2022; Hendra et al., 2022; Körtel et al., 2021; Meyer, 2019). We began with conventional MeRIP-seq to test feasibility in environmental samples using standard reagents and analysis, then can prioritize one of these low-input or antibody-free options in follow-up work if peak-level power is limited (Garcia-Campos et al., 2019; Ge et al., 2022; Hendra et al., 2022; Meyer, 2019). Low-input MeRIP refinements are also available if staying within an IP framework is preferred (Xia et al., 2024). For ultra-low m^6^A abundance, antibody-free/base-resolution chemistries and emerging biosensor concepts remain lab-prototype (Ahi & Khorshid, 2025; Khorshid & Ahi, 2025; Y. Yang et al., 2024), so they can be considered as future options rather than standard practice.

## Conclusion

In this study, we show that extra-organismal RNA shed by fish carries recoverable epitranscriptomic information and can be profiled with a conventional MeRIP-seq workflow. From tank water of *Labeotropheus trewavasae* exposed to 29 °C versus 26 °C, m^6^A-enriched eRNA libraries mapped reliably to the host (≈23% at stringent identity) and yielded substantially higher read mapping rates relative to comparable earlier studies, focusing on mRNA-seq approaches, potentially due to preferential m A enrichment in eukaryotic transcripts. We detected 412 differentially abundant eRNAs, including warmer-elevated signals in junction/adhesion and intermediate-filament programs and antigen processing, alongside warmer-decreased cohorts in glycolysis/PPP, nucleotide metabolism, and small-GTPase signaling; patterns aligned with thermal remodeling of barrier epithelia and metabolic re-set. Several important transcripts (e.g., *klf4, foxo3, slc1a5, ldha, mmp9, socs3*) have independent evidence of m^6^A regulation, supporting a mechanistic link between sustained warming and post-transcriptional control at water-exposed surfaces. Methodologically, correlations with filtered volume and biomass underscore the need to standardize capture effort and record covariates. While modest replication limited peak-level differential m^6^A inference and antibody-based assays warrant careful benchmarking, our results establish feasibility and provide the first evidence that environmental RNA retains m^6^A marks. Together, these findings position environmental epitranscriptomics as a practical extension of eRNA beyond transcript counts, enabling non-invasive, near-real-time readouts of physiological stress. Future work can pair optimized rRNA depletion with low-input or antibody-independent m^6^A methods and field validation, advancing m^6^A-guided biomonitoring of thermal stress in fishes and other aquatic taxa.

## Acknowledgements & Funding Information

We thank the Research Careers Campus Graz as well as the vice-rector for research of the University of Graz for financial support, enabling this study.

## References

Adiconis, X., Borges-Rivera, D., Satija, R., Deluca, D. S., Busby, M. A., Berlin, A. M., … Levin, J. Z. (2013). Comparative analysis of RNA sequencing methods for degraded or low-input samples. Nature Methods 2013 10:7, 10(7), 623–629. 10.1038/nmeth.2483

Ahi, E. P., & Khorshid, M. (2025). Potentials of RNA biosensors in developmental biology. Developmental Biology, 526, 173–188. 10.1016/J.YDBIO.2025.07.011

Ahi, E. P., Lindeza, A. S., Miettinen, A., & Primmer, C. R. (2025). Transcriptional responses to changing environments: insights from salmonids. Reviews in Fish Biology and Fisheries 2025, 1–26. 10.1007/S11160-025-09928-9

Ahi, E. P., & Schenekar, T. (2025). The Promise of Environmental RNA Research Beyond mRNA. Molecular Ecology, 34(12), e17787. 10.1111/MEC.17787

Ahi, E. P., & Singh, P. (2024). An emerging orchestrator of ecological adaptation: m6A regulation of posttranscriptional mechanisms. Molecular Ecology, 17545. 10.1111/MEC.17545

Akiyama, M. (2014). The roles of ABCA12 in epidermal lipid barrier formation and keratinocyte differentiation. Biochimica et Biophysica Acta (BBA) - Molecular and Cell Biology of Lipids, 1841(3), 435–440. 10.1016/J.BBALIP.2013.08.009

Andrews, S. (2012). FastQC: a quality control tool for high throughput sequence data. Retrieved from http://www.bioinformatics.babraham.ac.uk/projects/fastqc

Arroyo, J. D., Chevillet, J. R., Kroh, E. M., Ruf, I. K., Pritchard, C. C., Gibson, D. F., … Tewari, M. (2011). Argonaute2 complexes carry a population of circulating microRNAs independent of vesicles in human plasma. Proceedings of the National Academy of Sciences of the United States of America, 108(12), 5003–5008. 10.1073/PNAS.1019055108/SUPPL_FILE/ST02.DOC

Asakawa, S., Kraitavin, W., Yoshitake, K., Igarashi, Y., Mitsuyama, S., Kinoshita, S., … Watabe, S. (2019). Transcriptome Analysis of Yamame (Oncorhynchus masou) in Normal Conditions after Heat Stress. Biology 2019, Vol. 8, Page 21, 8(2), 21. 10.3390/BIOLOGY8020021

Basu, M., Paichha, M., Swain, B., Lenka, S. S., Singh, S., Chakrabarti, R., & Samanta, M. (2015). Modulation of TLR2, TLR4, TLR5, NOD1 and NOD2 receptor gene expressions and their downstream signaling molecules following thermal stress in the Indian major carp catla (Catla catla). 3 Biotech, 5(6), 1021–1030. 10.1007/S13205-015-0306-5/FIGURES/4

Bernardo, V. S., Torres, F. F., & da Silva, D. G. H. (2023). FoxO3 and oxidative stress: a multifaceted role in cellular adaptation. Journal of Molecular Medicine 2023 101:1, 101(1), 83–99. 10.1007/S00109-022-02281-5

Blackman, R., Couton, M., Keck, F., Kirschner, D., Carraro, L., Cereghetti, E., … Altermatt, F. (2024). Environmental DNA: The next chapter. Molecular Ecology, 33(11), e17355. 10.1111/MEC.17355

Bolger, A. M., Lohse, M., & Usadel, B. (2014). Trimmomatic: a flexible trimmer for Illumina sequence data. Bioinformatics, 30(15), 2114–2120. 10.1093/bioinformatics/btu170

Bowers, H. A., Pochon, X., Von Ammon, U., Gemmell, N., Stanton, J.-A. L., Jeunen, G.-J., … Towards, A. (2021). Towards the Optimization of eDNA/eRNA Sampling Technologies for Marine Biosecurity Surveillance. Water 2021, Vol. 13, Page 1113, 13(8), 1113. 10.3390/W13081113

Brawand, D., Wagner, C. E., Li, Y. I., Malinsky, M., Keller, I., Fan, S., … Di Palma, F. (2014). The genomic substrate for adaptive radiation in African cichlid fish. Nature, 513, 375–381. 10.1038/nature13726

Capo, E., Spong, G., Königsson, H., & Byström, P. (2020). Effects of filtration methods and water volume on the quantification of brown trout (Salmo trutta) and Arctic char (Salvelinus alpinus) eDNA concentrations via droplet digital PCR. Environmental DNA, 2(2), 152–160. 10.1002/EDN3.52

Chadzinska, M., Baginski, P., Kolaczkowska, E., Savelkoul, H. F. J., & Lidy Verburg-Van Kemenade, B. M. (2008). Expression profiles of matrix metalloproteinase 9 in teleost fish provide evidence for its active role in initiation and resolution of inflammation. Immunology, 125(4), 601–610. 10.1111/J.1365-2567.2008.02874.X

Chen, J., Wu, H., Zuo, T., Wu, J., & Chen, Z. (2025). METTL3⍰mediated N6⍰methyladenosine modification of MMP9 mRNA promotes colorectal cancer proliferation and migration. Oncology Reports, 53(1). 10.3892/OR.2024.8842

Cristescu, M. E. (2019). Can Environmental RNA Revolutionize Biodiversity Science? Trends in Ecology and Evolution, 34(8), 694–697. 10.1016/j.tree.2019.05.003

Dado-Senn, B., Skibiel, A. L., Dahl, G. E., Apelo, S. I. A., & Laporta, J. (2021). Dry period heat stress impacts mammary protein metabolism in the subsequent lactation. Animals, 11(9), 2676. 10.3390/ANI11092676/S1

Dokladny, K., Zuhl, M. N., & Moseley, P. L. (2016). Intestinal epithelial barrier function and tight junction proteins with heat and exercise. Journal of Applied Physiology, 120(6), 692–701. 10.1152/JAPPLPHYSIOL.00536.2015/ASSET/IMAGES/LARGE/ZDG9991516140003.JPEG

Dominissini, D., Moshitch-Moshkovitz, S., Schwartz, S., Salmon-Divon, M., Ungar, L., Osenberg, S., … Rechavi, G. (2012). Topology of the human and mouse m6A RNA methylomes revealed by m6A-seq. Nature 2012 485:7397, 485(7397), 201–206. 10.1038/nature11112

Duenser, A., Singh, P., Lecaudey, L. A., Sturmbauer, C., Albertson, R. C., Gessl, W., & Ahi, E. P. (2023). Conserved Molecular Players Involved in Human Nose Morphogenesis Underlie Evolution of the Exaggerated Snout Phenotype in Cichlids. Genome Biology and Evolution, 15(4). 10.1093/GBE/EVAD045

Fanidis, D., & Moulos, P. (2021). Integrative, normalization-insusceptible statistical analysis of RNA-Seq data, with improved differential expression and unbiased downstream functional analysis. Briefings in Bioinformatics, 22(3). 10.1093/BIB/BBAA156

Farrell, J. A., Whitmore, L., & Duffy, D. J. (2021). The Promise and Pitfalls of Environmental DNA and RNA Approaches for the Monitoring of Human and Animal Pathogens from Aquatic Sources. BioScience, 71(6), 609–625. 10.1093/BIOSCI/BIAB027

Fu, R., Jiang, X., Li, G., Zhu, Y., & Zhang, H. (2022). Junctional complexes in epithelial cells: sentinels for extracellular insults and intracellular homeostasis. The FEBS Journal, 289(23), 7314–7333. 10.1111/FEBS.16174

Fülle, J. B., de Almeida, R. A., Lawless, C., Stockdale, L., Yanes, B., Lane, E. B., … Ballestrem, C. (2024). Proximity Mapping of Desmosomes Reveals a Striking Shift in Their Molecular Neighborhood Associated With Maturation. Molecular & Cellular Proteomics, 23(3), 100735. 10.1016/J.MCPRO.2024.100735

Gann, E. R., Kang, Y., Dyhrman, S. T., Gobler, C. J., & Wilhelm, S. W. (2021). Metatranscriptome Library Preparation Influences Analyses of Viral Community Activity During a Brown Tide Bloom. Frontiers in Microbiology, 12, 664189. 10.3389/FMICB.2021.664189/BIBTEX

Gao, Z., Zha, X., Li, M., Xia, X., & Wang, S. (2024). Insights into the m6A demethylases FTO and ALKBH5⍰: structural, biological function, and inhibitor development. Cell & Bioscience 2024 14:1, 14(1), 1–20. 10.1186/S13578-024-01286-6

Garbo, S., D’Andrea, D., Colantoni, A., Fiorentino, F., Mai, A., Ramos, A., … Battistelli, C. (2024). m6A modification inhibits miRNAs’ intracellular function, favoring their extracellular export for intercellular communication. Cell Reports, 43(6), 114369. 10.1016/j.celrep.2024.114369

Garcia-Campos, M. A., Edelheit, S., Toth, U., Safra, M., Shachar, R., Viukov, S., … Schwartz, S. (2019). Deciphering the “m6A Code” via Antibody-Independent Quantitative Profiling. Cell, 178(3), 731-747.e16. 10.1016/J.CELL.2019.06.013/ATTACHMENT/B24F8FD4-A31B-47DB-B8E2-CC4730607C8C/MMC7.XLSX

Ge, R., Ye, C., Peng, Y., Dai, Q., Zhao, Y., Liu, S., … He, C. (2022). m6A-SAC-seq for quantitative whole transcriptome m6A profiling. Nature Protocols 2022 18:2, 18(2), 626–657. 10.1038/s41596-022-00765-9

Gifford, S. M., Sharma, S., Rinta-Kanto, J. M., & Moran, M. A. (2011). Quantitative analysis of a deeply sequenced marine microbial metatranscriptome. The ISME Journal, 5(3), 461–472. 10.1038/ISMEJ.2010.141

Giroux, M. S., Reichman, J. R., Langknecht, T., Burgess, R. M., & Ho, K. T. (2022). Environmental RNA as a Tool for Marine Community Biodiversity Assessments. Scientific Reports 2022 12:1, 12(1), 1–13. 10.1038/s41598-022-22198-w

Gleeson, J., Madugalle, S. U., Wan, C. Y., McLean, C., Bredy, T. W., De Paoli-Iseppi, R., & Clark, M. B. (2025). Isoform-level profiling of m6A epitranscriptomic signatures in human brain. Science Advances, 11(32), eadp0783. 10.1126/SCIADV.ADP0783/SUPPL_FILE/SCIADV.ADP0783_TABLES_S3_TO_S15.ZIP

Gross, A., Pack, L. A. P., Schacht, G. M., Kant, S., Ungewiss, H., Meir, M., … Strnad, P. (2018). Desmoglein 2, but not desmocollin 2, protects intestinal epithelia from injury. Mucosal Immunology, 11(6), 1630–1639. 10.1038/S41385-018-0062-Z

Grossi, V., Forte, G., Sanese, P., Peserico, A., Tezil, T., Signorile, M. L., … Simone, C. (2018). The longevity SNP rs2802292 uncovered: HSF1 activates stress-dependent expression of FOXO3 through an intronic enhancer. Nucleic Acids Research, 46(11), 5587. 10.1093/NAR/GKY331

Guo, C., Hu, Y., Cao, X., & Wang, Y. (2021). HILIC-MS/MS for the Determination of Methylated Adenine Nucleosides in Human Urine. Analytical Chemistry, 93(51), 17060–17068. 10.1021/ACS.ANALCHEM.1C03829/SUPPL_FILE/AC1C03829_SI_001.PDF

Hao, W. C., Dian, M. J., Zhou, Y., Zhong, Q. L., Pang, W. Q., Li, Z. J., … Xiao, D. (2022). Autophagy induction promoted by m6A reader YTHDF3 through translation upregulation of FOXO3 mRNA. Nature Communications 2022 13:1, 13(1), 1–23. 10.1038/s41467-022-32963-0

Hechler, R. M., Yates, M. C., Chain, F. J. J., & Cristescu, M. E. (2023). Environmental transcriptomics under heat stress: Can environmental RNA reveal changes in gene expression of aquatic organisms?, 1–15. 10.1111/MEC.17152

Hendra, C., Pratanwanich, P. N., Wan, Y. K., Goh, W. S. S., Thiery, A., & Göke, J. (2022). Detection of m6A from direct RNA sequencing using a multiple instance learning framework. Nature Methods, 19(12), 1590–1598. 10.1038/S41592-022-01666-1

Herbert, Z. T., Kershner, J. P., Butty, V. L., Thimmapuram, J., Choudhari, S., Alekseyev, Y. O., … Levine, S. S. (2018). Cross-site comparison of ribosomal depletion kits for Illumina RNAseq library construction. BMC Genomics, 19(1), 1–10. 10.1186/S12864-018-4585-1/TABLES/1

Hiki, K., Watanabe, H., & Yamamoto, H. (2024). Relative gene expression analysis of catalase in environmental RNA from Japanese medaka exposed to toxic chemicals. Environmental DNA, 6(2), e532. 10.1002/EDN3.532

Hiki, K., Yamagishi, T., & Yamamoto, H. (2023). Environmental RNA as a Noninvasive Tool for Assessing Toxic Effects in Fish: A Proof-of-concept Study Using Japanese Medaka Exposed to Pyrene. Environmental Science and Technology, 57(34), 12654–12662. 10.1021/ACS.EST.3C03737/SUPPL_FILE/ES3C03737_SI_001.XLSX

Hinlo, R., Gleeson, D., Lintermans, M., & Furlan, E. (2017). Methods to maximise recovery of environmental DNA from water samples. PLOS ONE, 12(6), e0179251. 10.1371/JOURNAL.PONE.0179251

Hu, Y., Fang, Z., Mu, J., Huang, Y., Zheng, S., Yuan, Y., & Guo, C. (2021). Quantitative Analysis of Methylated Adenosine Modifications Revealed Increased Levels of N6-Methyladenosine (m6A) and N6,2⍰-O-Dimethyladenosine (m6Am) in Serum From Colorectal Cancer and Gastric Cancer Patients. Frontiers in Cell and Developmental Biology, 9, 694673. 10.3389/FCELL.2021.694673/BIBTEX

Johnson, J. L., Najor, N. A., & Green, K. J. (2014). Desmosomes: Regulators of Cellular Signaling and Adhesion in Epidermal Health and Disease. Cold Spring Harbor Perspectives in Medicine, 4(11), a015297. 10.1101/CSHPERSPECT.A015297

Kagzi, K., Millette, K. L., Littlefair, J. E., Pochon, X., Wood, S. A., Fussmann, G. F., & Cristescu, M. E. (2023). Assessing the degradation of environmental DNA and RNA based on genomic origin in a metabarcoding context. Environmental DNA, 5(5), 1016–1031. 10.1002/EDN3.437

Kayser, J., Haslbeck, M., Dempfle, L., Krause, M., Grashoff, C., Buchner, J., … Bausch, A. R. (2013). The small heat shock protein Hsp27 affects assembly dynamics and structure of keratin intermediate filament networks. Biophysical Journal, 105(8), 1778–1785. 10.1016/j.bpj.2013.09.007

Keen, A. N., Mackrill, J. J., Gardner, P., & Shiels, H. A. (2021). Compliance of the fish outflow tract is altered by thermal acclimation through connective tissue remodelling. Journal of the Royal Society Interface, 18(184). 10.1098/RSIF.2021.0492

Khorshid, M., & Ahi, E. P. (2025). RNA Aptamers and Epitranscriptomics: Charting Unexplored Territories in RNA Biology. 10.20944/PREPRINTS202509.1838.V1

Köberle, V., Pleli, T., Schmithals, C., Augusto Alonso, E., Haupenthal, J., Bönig, H., … Piiper, A. (2013). Differential Stability of Cell-Free Circulating microRNAs: Implications for Their Utilization as Biomarkers. PLOS ONE, 8(9), e75184. 10.1371/JOURNAL.PONE.0075184

Körtel, N., Rücklé, C., Zhou, Y., Busch, A., Hoch-Kraft, P., Sutandy, F. X. R., … Zarnack, K. (2021). Deep and accurate detection of m6A RNA modifications using miCLIP2 and m6Aboost machine learning. Nucleic Acids Research, 49(16), e92–e92. 10.1093/NAR/GKAB485

Langmead, B., & Salzberg, S. L. (2012). Fast gapped-read alignment with Bowtie 2. Nature Methods 2012 9:4, 9(4), 357–359. 10.1038/nmeth.1923

Leprince, C., & Simon, M. (2025). Epidermal lamellar bodies, essential organelles for the skin barrier. Frontiers in Cell and Developmental Biology, 13, 1597884. 10.3389/FCELL.2025.1597884/BIBTEX

Li, H., & Durbin, R. (2009). Fast and accurate short read alignment with Burrows–Wheeler transform. Bioinformatics, 25(14), 1754–1760. 10.1093/BIOINFORMATICS/BTP324

Li, Q., Ni, Y., Zhang, L., Jiang, R., Xu, J., Yang, H., … Wang, X. (2021). HIF-1α-induced expression of m6A reader YTHDF1 drives hypoxia-induced autophagy and malignancy of hepatocellular carcinoma by promoting ATG2A and ATG14 translation. Signal Transduction and Targeted Therapy 2021 6:1, 6(1), 1–13. 10.1038/s41392-020-00453-8

Li, Y., Zheng, Q., Bao, C., Li, S., Guo, W., Zhao, J., … Huang, S. (2015). Circular RNA is enriched and stable in exosomes: a promising biomarker for cancer diagnosis. Cell Research 2015 25:8, 25(8), 981–984. 10.1038/cr.2015.82

Li, Z., Meng, X., Chen, Y., Xu, X., & Guo, J. (2023). N6-methyladenosine (m6A) writer METTL3 accelerates the apoptosis of vascular endothelial cells in high glucose. Heliyon, 9(3), e13721. 10.1016/j.heliyon.2023.e13721

Lin, C., Zeng, M., Song, J., Li, H., Feng, Z., Li, K., & Pei, Y. (2023). PRRSV alters m6A methylation and alternative splicing to regulate immune, extracellular matrix-associated function. International Journal of Biological Macromolecules, 253, 126741. 10.1016/J.IJBIOMAC.2023.126741

Lin, T., & Meegaskumbura, M. (2025). Fish MicroRNA Responses to Thermal Stress: Insights and Implications for Aquaculture and Conservation Amid Global Warming. Animals, 15(5), 624. 10.3390/ANI15050624/S1

Littlefair, J. E., Rennie, M. D., & Cristescu, M. E. (2022). Environmental nucleic acids: A field-based comparison for monitoring freshwater habitats using eDNA and eRNA. Molecular Ecology Resources, 22(8), 2928. 10.1111/1755-0998.13671

Liu, Y., Wang, J., Yi, Y., Zhang, H., Liu, J., Liu, M., … Xiao, X. (2006). Induction of KLF4 in response to heat stress. Cell Stress & Chaperones, 11(4), 379. 10.1379/CSC-210.1

Ma, L., Xue, X., Zhang, X., Yu, K., Xu, X., Tian, X., … Wang, J. (2022). The essential roles of m6A RNA modification to stimulate ENO1-dependent glycolysis and tumorigenesis in lung adenocarcinoma. Journal of Experimental and Clinical Cancer Research, 41(1), 1–21. 10.1186/S13046-021-02200-5/FIGURES/8

Macher, T. H., Arle, J., Beermann, A. J., Frank, L., Hupało, K., Koschorreck, J., … Leese, F. (2024). Is it worth the extra mile? Comparing environmental DNA and RNA metabarcoding for vertebrate and invertebrate biodiversity surveys in a lowland stream. PeerJ, 12(10), e18016. 10.7717/PEERJ.18016/SUPP-15

Madeira, D., Vinagre, C., & Diniz, M. S. (2016). Are fish in hot water? Effects of warming on oxidative stress metabolism in the commercial species Sparus aurata. Ecological Indicators, 63, 324–331. 10.1016/J.ECOLIND.2015.12.008

Matsuda, M., Chu, C. W., & Sokol, S. Y. (2022). Lmo7 recruits myosin II heavy chain to regulate actomyosin contractility and apical domain size in Xenopus ectoderm. Development (Cambridge), 149(10). 10.1242/DEV.200236/VIDEO-1

McIntyre, A. B. R., Gokhale, N. S., Cerchietti, L., Jaffrey, S. R., Horner, S. M., & Mason, C. E. (2020). Limits in the detection of m6A changes using MeRIP/m6A-seq. Scientific Reports 2020 10:1, 10(1), 1–15. 10.1038/s41598-020-63355-3

Mehravar, M., Kumar, Y., Olshansky, M., Dakle, P., Bullen, M., Pandey, V. K., … Das, P. P. (2022). MOV10 facilitates messenger RNA decay in an N6-methyladenosine (m6A) dependent manner to maintain the mouse embryonic stem cells state. BioRxiv, 2021.08.11.456030. 10.1101/2021.08.11.456030

Mei, Z., Mou, Y., Zhang, N., Liu, X., He, Z., & Gu, S. (2023). Emerging Mutual Regulatory Roles between m6A Modification and microRNAs. International Journal of Molecular Sciences, 24(1). 10.3390/IJMS24010773

Meyer, K. D. (2019). DART-seq: an antibody-free method for global m6A detection. Nature Methods 2019 16:12, 16(12), 1275–1280. 10.1038/s41592-019-0570-0

Miyata, K., Inoue, Y., Amano, Y., Nishioka, T., Yamane, M., Kawaguchi, T., … Honda, H. (2021). Fish environmental RNA enables precise ecological surveys with high positive predictivity. Ecological Indicators, 128, 107796. 10.1016/J.ECOLIND.2021.107796

Miyata, K., Inoue, Y., Yamane, M., & Honda, H. (2025). Fish environmental RNA sequencing sensitively captures accumulative stress responses through short-term aquarium sampling. Science of The Total Environment, 959, 178182. 10.1016/J.SCITOTENV.2024.178182

Moulos, P., & Hatzis, P. (2015). Systematic integration of RNA-Seq statistical algorithms for accurate detection of differential gene expression patterns. Nucleic Acids Research, 43(4), e25–e25. 10.1093/NAR/GKU1273

Murshid, A., Gong, J., & Calderwood, S. K. (2012). The role of heat shock proteins in antigen cross presentation. Frontiers in Immunology, 3(MAR), 20610. 10.3389/FIMMU.2012.00063/BIBTEX

Ooshio, T., Irie, K., Morimoto, K., Fukuhara, A., Imai, T., & Takai, Y. (2004). Involvement of LMO7 in the Association of Two Cell-Cell Adhesion Molecules, Nectin and E-cadherin, through Afadin and α-Actinin in Epithelial Cells. Journal of Biological Chemistry, 279(30), 31365–31373. 10.1074/JBC.M401957200

Pan, H. Y., Yu, Y., Cao, T., Liu, Y., Zhou, Y. L., & Zhang, X. X. (2021). Systematic Profiling of Exosomal Small RNA Epigenetic Modifications by High-Performance Liquid Chromatography-Mass Spectrometry. Analytical Chemistry, 93(45), 14907–14911. 10.1021/ACS.ANALCHEM.1C03869/ASSET/IMAGES/LARGE/AC1C03869_0004.JPEG

Patra, A. K., & Kar, I. (2021). Heat stress on microbiota composition, barrier integrity, and nutrient transport in gut, production performance, and its amelioration in farm animals. Journal of Animal Science and Technology, 63(2), 211. 10.5187/JAST.2021.E48

Pereira-Marques, J., Ferreira, R. M., & Figueiredo, C. (2024). A metatranscriptomics strategy for efficient characterization of the microbiome in human tissues with low microbial biomass. Gut Microbes, 16(1). 10.1080/19490976.2024.2323235

Pochon, X., Bowers, H. A., Zaiko, A., & Wood, S. A. (2025). Advancing the environmental DNA and RNA toolkit for aquatic ecosystem monitoring and management. PeerJ, 13(3), e19119. 10.7717/PEERJ.19119

Pu, X., Wu, Y., Long, W., Sun, X., Yuan, X., Wang, D., … Xu, M. (2025). The m6A reader IGF2BP2 promotes pancreatic cancer progression through the m6A-SLC1A5-mTORC1 axis. Cancer Cell International, 25(1), 1–15. 10.1186/S12935-025-03736-8/FIGURES/5

Qin, H., Long, Z., Huang, Z., Ma, J., Kong, L., Lin, Y., … Li, Z. (2023). A Comparison of the Physiological Responses to Heat Stress of Two Sizes of Juvenile Spotted Seabass (Lateolabrax maculatus). Fishes 2023, Vol. 8, Page 340, 8(7), 340. 10.3390/FISHES8070340

Qin, S., Zhang, Q., Xu, Y., Ma, S., Wang, T., Huang, Y., & Ju, S. (2022). m6A-modified circRNAs: detections, mechanisms, and prospects in cancers. Molecular Medicine 2022 28:1, 28(1), 1–17. 10.1186/S10020-022-00505-5

Quan, J., Kang, Y., Luo, Z., Zhao, G., Li, L., & Liu, Z. (2021). Integrated analysis of the responses of a circRNA-miRNA-mRNA ceRNA network to heat stress in rainbow trout (Oncorhynchus mykiss) liver. BMC Genomics, 22(1), 1–10. 10.1186/S12864-020-07335-X/FIGURES/5

R Core Team. (2023). R: A Language and Environment for Statistical Computing. R Foundation for Statistical Computing. Vienna, Austria. 10.1007/978-3-540-74686-7

Roberts, R. B., Hu, Y., Albertson, R. C., & Kocher, T. D. (2011). Craniofacial divergence and ongoing adaptation via the hedgehog pathway. Proceedings of the National Academy of Sciences of the United States of America, 108(32), 13194–13199. 10.1073/pnas.1018456108

Seo, H., Cho, Y. C., Ju, A., Lee, S., Park, B. C., Park, S. G., … Cho, S. (2017). Dual-specificity phosphatase 5 acts as an anti-inflammatory regulator by inhibiting the ERK and NF-κB signaling pathways. Scientific Reports 2017 7:1, 7(1), 1–12. 10.1038/s41598-017-17591-9

Shen, C., Xuan, B., Yan, T., Ma, Y., Xu, P., Tian, X., … Hong, J. (2020). M6A-dependent glycolysis enhances colorectal cancer progression. Molecular Cancer, 19(1), 1–19. 10.1186/S12943-020-01190-W/FIGURES/7

Silva, S. S., Lopes, C., Teixeira, A. L., Sousa, M. J. C. De, & Medeiros, R. (2015). Forensic miRNA: Potential biomarker for body fluids? Forensic Science International: Genetics, 14, 1–10. 10.1016/J.FSIGEN.2014.09.002

Spens, J., Evans, A. R., Halfmaerten, D., Knudsen, S. W., Sengupta, M. E., Mak, S. S. T., … Hellström, M. (2017). Comparison of capture and storage methods for aqueous macrobial eDNA using an optimized extraction protocol: advantage of enclosed filter. Methods in Ecology and Evolution, 8(5), 635–645. 10.1111/2041-210X.12683

Stevens, J. D., & Parsley, M. B. (2023). Environmental RNA applications and their associated gene targets for management and conservation. Environmental DNA, 5(2), 227–239. 10.1002/EDN3.386

Stincone, A., Prigione, A., Cramer, T., Wamelink, M. M. C., Campbell, K., Cheung, E., … Ralser, M. (2015). The return of metabolism: biochemistry and physiology of the pentose phosphate pathway. Biological Reviews, 90(3), 927–963. 10.1111/BRV.12140

Sun, Y., Sun, Y., He, X., Li, S., Xu, X., Feng, Y., … Sun, G. (2024). Transcriptome-wide methylated RNA immunoprecipitation sequencing profiling reveals m6A modification involved in response to heat stress in Apostichopus japonicus. BMC Genomics, 25(1), 1071. 10.1186/S12864-024-10972-1/FIGURES/4

Toivola, D. M., Strnad, P., Habtezion, A., & Omary, M. B. (2010). Intermediate filaments take the heat as stress proteins. Trends in Cell Biology, 20(2), 79–91. 10.1016/J.TCB.2009.11.004

Veilleux, H. D., Misutka, M. D., & Glover, C. N. (2021). Environmental DNA and environmental RNA: Current and prospective applications for biological monitoring. Science of The Total Environment, 782, 146891. 10.1016/J.SCITOTENV.2021.146891

Wang, Q., Zhang, S., He, X., Li, S., Xu, X., Feng, Y., … Sun, G. (2025). Integrated m6A RNA methylation and transcriptomic analysis of Apostichopus japonicus under combined high-temperature and hypoxia stress. BMC Genomics, 26(1), 363. 10.1186/S12864-025-11532-X

Wang, S., Zhang, K., Tan, S., Xin, J., Yuan, Q., Xu, H., … Du, M. (2021). Circular RNAs in body fluids as cancer biomarkers: the new frontier of liquid biopsies. Molecular Cancer, 20(1), 1–10. 10.1186/S12943-020-01298-Z/METRICS

Wang, Yong, Yang, X., Li, S., Wu, Q., Guo, H., Wang, H., … Wang, J. (2023). Research Note: Heat stress affects immune and oxidative stress indices of the immune organs of broilers by changing the expressions of adenosine triphosphate-binding cassette subfamily G member 2, sodium-dependent vitamin C transporter-2, and mitochondrial calcium uniporter. Poultry Science, 102(8), 102814. 10.1016/J.PSJ.2023.102814

Wang, Yue, Su, C., Liu, Q., Hao, X., Han, S., Doretto, L. B., … Wang, Q. (2024). Transcriptome Analysis Revealed the Early Heat Stress Response in the Brain of Chinese Tongue Sole (Cynoglossus semilaevis). Animals, 14(1), 84. 10.3390/ANI14010084/S1

Wingett, S. W., & Andrews, S. (2018). FastQ Screen: A tool for multi-genome mapping and quality control. F1000Research, 7, 1338. 10.12688/F1000RESEARCH.15931.2

Wood, S. A., Biessy, L., Latchford, J. L., Zaiko, A., von Ammon, U., Audrezet, F., … Pochon, X. (2020). Release and degradation of environmental DNA and RNA in a marine system. Science of The Total Environment, 704, 135314. 10.1016/J.SCITOTENV.2019.135314

Xia, W., Guo, L., Su, H., Li, J., Lu, J., Li, H., & Huang, B. (2024). A low-cost, low-input method establishment for m6A MeRIP-seq. Bioscience Reports, 44(1), 20231430. 10.1042/BSR20231430/233871

Yang, S., Li, D., Feng, L., Zhang, C., Xi, D., Liu, H., … Du, X. (2023). Transcriptome analysis reveals the high temperature induced damage is a significant factor affecting the osmotic function of gill tissue in Siberian sturgeon (Acipenser baerii). BMC Genomics, 24(1), 1–12. 10.1186/S12864-022-08969-9/TABLES/2

Yang, Y., Lu, Y., Wang, Y., Wen, X., Qi, C., Piao, W., & Jin, H. (2024). Current progress in strategies to profile transcriptomic m6A modifications. Frontiers in Cell and Developmental Biology, 12, 1392159. 10.3389/FCELL.2024.1392159/XML

Yates, M. C., Furlan, E., Thalinger, B., Yamanaka, H., & Bernatchez, L. (2023). Beyond species detection— leveraging environmental DNA and environmental RNA to push beyond presence/absence applications. Environmental DNA, 5(5), 829–835. 10.1002/EDN3.459

Yates, Matthew C., Derry, A. M., & Cristescu, M. E. (2021). Environmental RNA: A Revolution in Ecological Resolution? Trends in Ecology and Evolution, 36(7), 601–609. 10.1016/j.tree.2021.03.001

Zaccara, S., Ries, R. J., & Jaffrey, S. R. (2019). Reading, writing and erasing mRNA methylation. Nature Reviews Molecular Cell Biology, 20(10), 608–624. 10.1038/s41580-019-0168-5

Zhang, K., Zhang, T., Yang, Y., Tu, W., Huang, H., Wang, Y., … Chen, Z. (2022). N6-methyladenosine-mediated LDHA induction potentiates chemoresistance of colorectal cancer cells through metabolic reprogramming. Theranostics, 12(10), 4802. 10.7150/THNO.73746

Zhang, Y., Qiu, Y., Liu, K., Zhong, W., Yang, J., Altermatt, F., & Zhang, X. (2024). Evaluating eDNA and eRNA metabarcoding for aquatic biodiversity assessment: From bacteria to vertebrates. Environmental Science and Ecotechnology, 21, 100441. 10.1016/J.ESE.2024.100441

Zhou, C., Gao, P., & Wang, J. (2023). Comprehensive Analysis of Microbiome, Metabolome, and Transcriptome Revealed the Mechanisms of Intestinal Injury in Rainbow Trout under Heat Stress. International Journal of Molecular Sciences, 24(10), 8569. 10.3390/IJMS24108569/S1

Zhu, X., Wang, J., Zhang, H., Yue, H., Zhu, J., Li, J., … Shi, T. (2024). Downregulated KLF4, induced by m6A modification, aggravates intestinal barrier dysfunction in inflammatory bowel disease. Cellular and Molecular Life Sciences, 81(1), 1–23. 10.1007/S00018-024-05514-7/FIGURES/9

Ahi, E. P., & Singh, P. (2024). An emerging orchestrator of ecological adaptation: m6A regulation of post-transcriptional mechanisms. Molecular Ecology, 17545. 10.1111/MEC.17545

Garcia-Campos, M. A., Edelheit, S., Toth, U., Safra, M., Shachar, R., Viukov, S., … Schwartz, S. (2019). Deciphering the “m6A Code” via Antibody-Independent Quantitative Profiling. Cell, 178(3), 731-747.e16. 10.1016/J.CELL.2019.06.013/ATTACHMENT/B24F8FD4-A31B-47DB-B8E2-C4730607C8C/MMC7.XLSX

Hechler, R. M., Yates, M. C., Chain, F. J. J., & Cristescu, M. E. (2023). Environmental transcriptomics under heat stress: Can environmental RNA reveal changes in gene expression of aquatic organisms? Molecular Ecology, 00, 1–15. 10.1111/MEC.17152

Zhang, K., Zhang, T., Yang, Y., Tu, W., Huang, H., Wang, Y., … Chen, Z. (2022). N6-methyladenosine-mediated LDHA induction potentiates chemoresistance of colorectal cancer cells through metabolic reprogramming. Theranostics, 12(10), 4802. 10.7150/THNO.73746

